# Broad-spectrum adaptive antibiotic resistance associated with *Pseudomonas aeruginosa* mucin-dependent surfing motility

**DOI:** 10.1101/309625

**Authors:** Evelyn Sun, Erin E. Gill, Reza Falsafi, Amy Yeung, Sijie Liu, Robert E.W. Hancock

## Abstract

Surfing motility is a novel form of surface adaptation exhibited by the nosocomial pathogen, *Pseudomonas aeruginosa*, in the presence of the glycoprotein mucin that is found in high abundance at mucosal surfaces especially the lungs of cystic fibrosis and bronchiectasis patients. Here we investigated the adaptive antibiotic resistance of *P. aeruginosa* under conditions in which surfing occurs compared to cells undergoing swimming. *P. aeruginosa* surfing cells were significantly more resistant to several classes of antibiotics including aminoglycosides, carbapenems, polymyxins, and fluroquinolones. This was confirmed by incorporation of antibiotics into growth medium, which revealed a concentration-dependent inhibition of surfing motility that occurred at concentrations much higher than those needed to inhibit swimming. To investigate the basis of resistance, RNA-Seq was performed and revealed that surfing influenced the expression of numerous genes. Included amongst genes dysregulated under surfing conditions were multiple genes from the *Pseudomonas* resistome, which are known to affect antibiotic resistance when mutated. Screening transposon mutants in these surfing-dysregulated resistome genes revealed that several of these mutants exhibited changes in susceptibility to one or more antibiotics under surfing conditions, consistent with a contribution to the observed adaptive resistance. In particular, several mutants in resistome genes, including *armR, recG, atpB, clpS, nuoB*, and certain hypothetical genes such as PA5130, PA3576 and PA4292, showed contributions to broad-spectrum resistance under surfing conditions and could be complemented by their respective cloned genes. Therefore, we propose that surfing adaption led to extensive multidrug adaptive resistance as a result of the collective dysregulation of diverse genes.

## Introduction

The rise of antibiotic resistance is a global concern. As the number of new antibiotics being discovered declines and the extensive and sometimes inappropriate use of antibiotics continues, more patients suffer and die from infections caused by antibiotic resistant bacteria (1, 2). *Pseudomonas aeruginosa* is categorized as a serious threat by the CDC and a priority 1 critical pathogen by the World Health Organization (WHO) (3–5). *P. aeruginosa* is a Gram-negative opportunistic pathogen that causes approximately 10% of all hospital-acquired infections as well as chronic infections in the lungs of individuals with cystic fibrosis (CF) (6, 7). Therefore, there is a growing need to understand the mechanisms leading to antibiotic resistance in *P. aeruginosa*.

*P. aeruginosa* can deploy intrinsic, acquired and adaptive resistance mechanisms (8, 9). Intrinsic resistance refers to the bacterium’s natural qualities that allow it to evade the effects of antibiotics and is not induced by stress nor the presence of antibiotics (8). Intrinsic resistance includes low outer membrane permeability that works in synergy with intrinsic levels of efflux pumps, β-lactamases, periplasmic enzymes, and other resistance mechanisms. Acquired resistance can be selected for by antibiotic exposure and occurs due to mutations or the acquisition of genetic elements such as plasmids, transposons, and integrons (8, 9). Adaptive resistance refers to resistance that occurs due to environmental circumstances and is thought to be largely due to transcriptional changes in genes that determine resistance/susceptibility and is reversible when environmental circumstances (e.g. exposure to stresses including antibiotics, complex adaptive growth states such as swarming or biofilm formation, etc.) are reversed (8). *P. aeruginosa* in the cystic fibrosis lung, especially during late stages, is thought to grow as biofilms (10). Biofilms represent a growth state in which bacteria grow as structured communities on surfaces and exhibit multidrug adaptive resistance (11).

Bacterial motility is a critical aspect of *P. aeruginosa* pathogenesis. Motility is needed for colonization of the host and the establishment of biofilms (12). It is also often coupled with the expression of virulence factors. *P. aeruginosa* has two known forms of polar locomotory appendages, a tail-like flagellum and hair-like type IV pilus; these contribute to a diverse set of motile phenotypes or lifestyles. The three most highly-studied forms of motility are swimming, twitching and swarming (12, 13). Swimming motility involves the use of flagellar rotation to move within aqueous environments. Twitching depends on the type IV pilus to enable movement on solid surfaces through the extension and retraction of polar pili (12, 14). These types of movements are not accompanied by major changes in gene expression. In contrast, swarming motility is a complex adaptation that involves multi-cellular coordination to enable movement on semi-solid surfaces in the presence of a poor nitrogen source (12, 14). This surface motility form is dependent on both pili and flagella, and results in dendritic (*P. aeruginosa* strain PA14) or solar flare colonies (*P. aeruginosa* strain PA01) on 0.4-0.6% (wt/vol) agar (12). A less studied form of motility termed sliding occurs when *Pseudomonas* glide on solid surfaces independent of any appendages but dependent on the production of rhamnolipid surfactants to reduce surface tension (15). The conditions under which swarming motility occur have been proposed to reflect CF lung conditions due to the high similarities in composition (semi viscous surface, amino acids as a nitrogen source, glucose as a carbon source); however, swarming models lack a major glycoprotein known as mucin found in the CF lungs and involved in regulating mucosal viscosity (12). By incorporating mucin into an artificial CF model, a novel form of motility termed “surfing” was discovered (12).

Mucin is secreted from mucosal and submucosal glands in the lungs and other mucosal surfaces (12). It contains a polypeptide core with branched oligosaccharide chains. Molecular cross-linking of its structure contributes to the viscoelastic properties of mucus. When mucin is added to media that normally support swimming or swarming, accelerated surface motility termed surfing occurs. Surfing depends on intact flagella but not type IV pili (12). Surfing colonies appear relatively circular with thick white outer edges containing mostly non-flagellated cells piled on top of each other and a blue-green centre with flagellated cells (12). Unlike swarming, surfing motility does not require such strict growth conditions and can occur in nutrient-rich or minimal medium, in the presence of ammonium as a nitrogen source and at a range of viscosities/agar concentrations (ranging from 0.3% to 1.0% wt/vol). Mucin is proposed to act as a wetting agent or lubricant and, unlike swarming or sliding, surfing does not depend on rhamnolipid production (12). It was suggested that surfing is a complex adaptive form of motility (12).

Here we demonstrate that surfing cells exhibited multi-drug adaptive resistance, dependent on the complex adaptive changes that accompanied this motility phenotype. Compared to swimming, surfing adaptive cells were significantly more resistant to several classes of antibiotics including aminoglycosides, polymyxins, flouroquinolones, and carbapenems. Screening mutants of resistome genes that were found to be dysregulated under surfing conditions revealed changes in susceptibility that may account for their contribution to the observed resistance.

## Results

### Surfing cells exhibited broad-spectrum antibiotic resistance

Disk diffusion assay results (Fig 1), assessing how close surfing and swimming cells approached an antibiotic disk, revealed a significant decrease in the zone of inhibition under surfing conditions (SCFM 0.3% agar, 0.4% mucin) when compared to swimming (SCFM 0.3% agar) or to a disk diffusion lawn control/growth control (SCFM 1.5% agar). This was observed for 12 of the 17 antibiotics tested with the exceptions of 3 of the β-lactams and 2 macrolides. Compared to swimming bacteria (and disk diffusion assays), surfing cells exhibited significant adaptive resistance to the tested aminoglycosides, carbapenems, polymyxins, fluoroquinolones, trimethoprim, tetracycline, and chloramphenicol, with complete resistance to 3 different aminoglycosides, imipenem, clarithromycin, and the polymyxins.

**Fig. 1.**
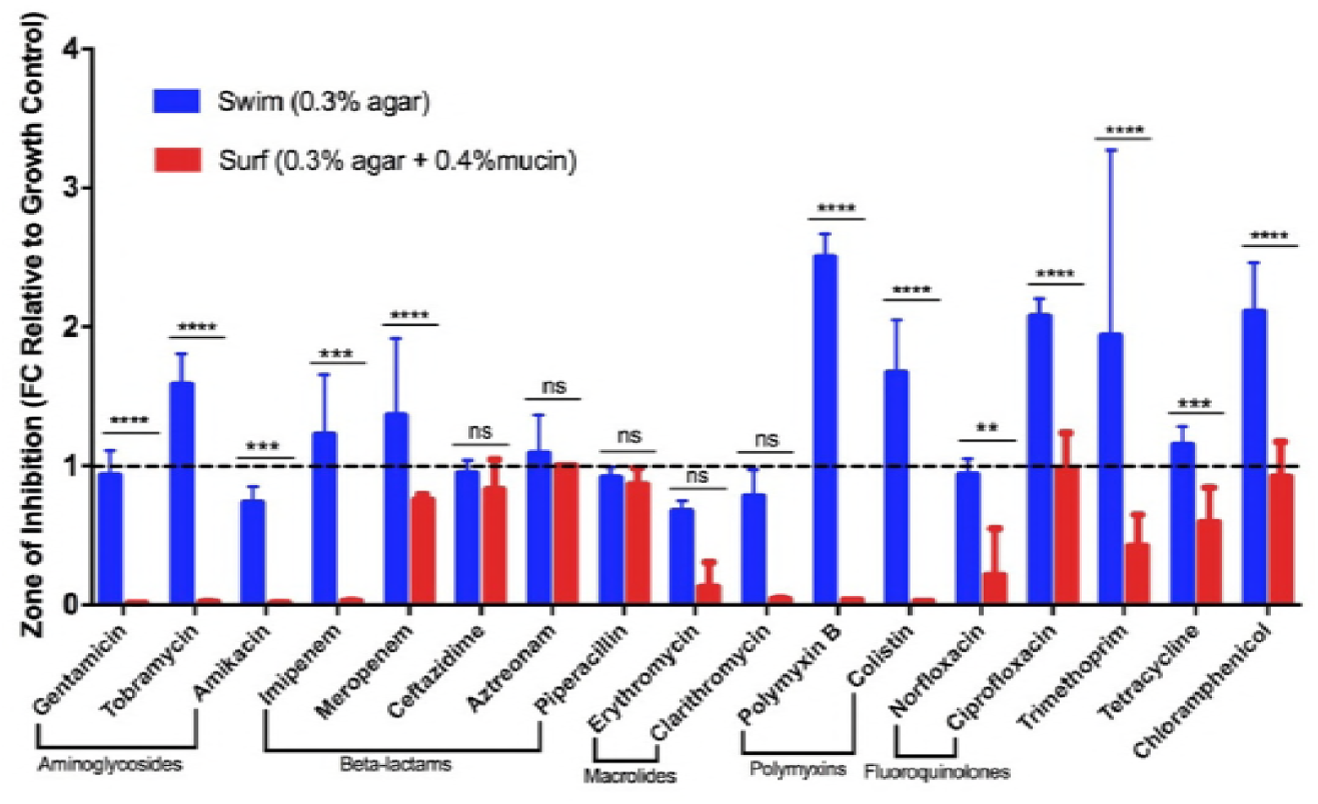
Multi-drug adaptive resistance of surfing colonies. Fold change of the zone of inhibition under swimming (0.3% agar) and surfing (0.3% agar 0.4% mucin) conditions relative to a disk diffusion control (lawn plated on 1.5% agar without mucin) in MSCFM which is set as 1 as indicated by the dashed line. Statistical significance between swimming and surfing as determined using two-way ANOVA. (n=3). * p<0.5, * * p<0.01, * * * p< 10^-3^, * * * * p<10^-4^.

### Antibiotic incorporation assays to confirm adaptive resistance

To further investigate the adaptive resistance of surfing colonies, 5 selected antibiotics were incorporated into growth plates to determine how they affect the initiation and propagation of motility colonies. These antibiotic incorporation assays (Fig 2), involving norfloxacin, polymyxin B, imipenem, tetracycline, and tobramycin, revealed a concentration-dependent inhibition of surfing motility and showed that surfing motility proceeded at antibiotic concentrations that completely inhibited swimming. For example, surfing occurred on 0.1 μM imipenem whereas swimming was completely abolished at this concentration. As the imipenem concentration increased, there was a clear reduction in the size of the surfing colony and at a concentration of 1 μM imipenem both surfing and swimming were completely inhibited. Indeed, for all five antibiotics tested, inhibition of surfing occurred with increasing concentrations but still occurred to some extent at concentrations much higher than those inhibiting swimming.

**Fig. 2.**
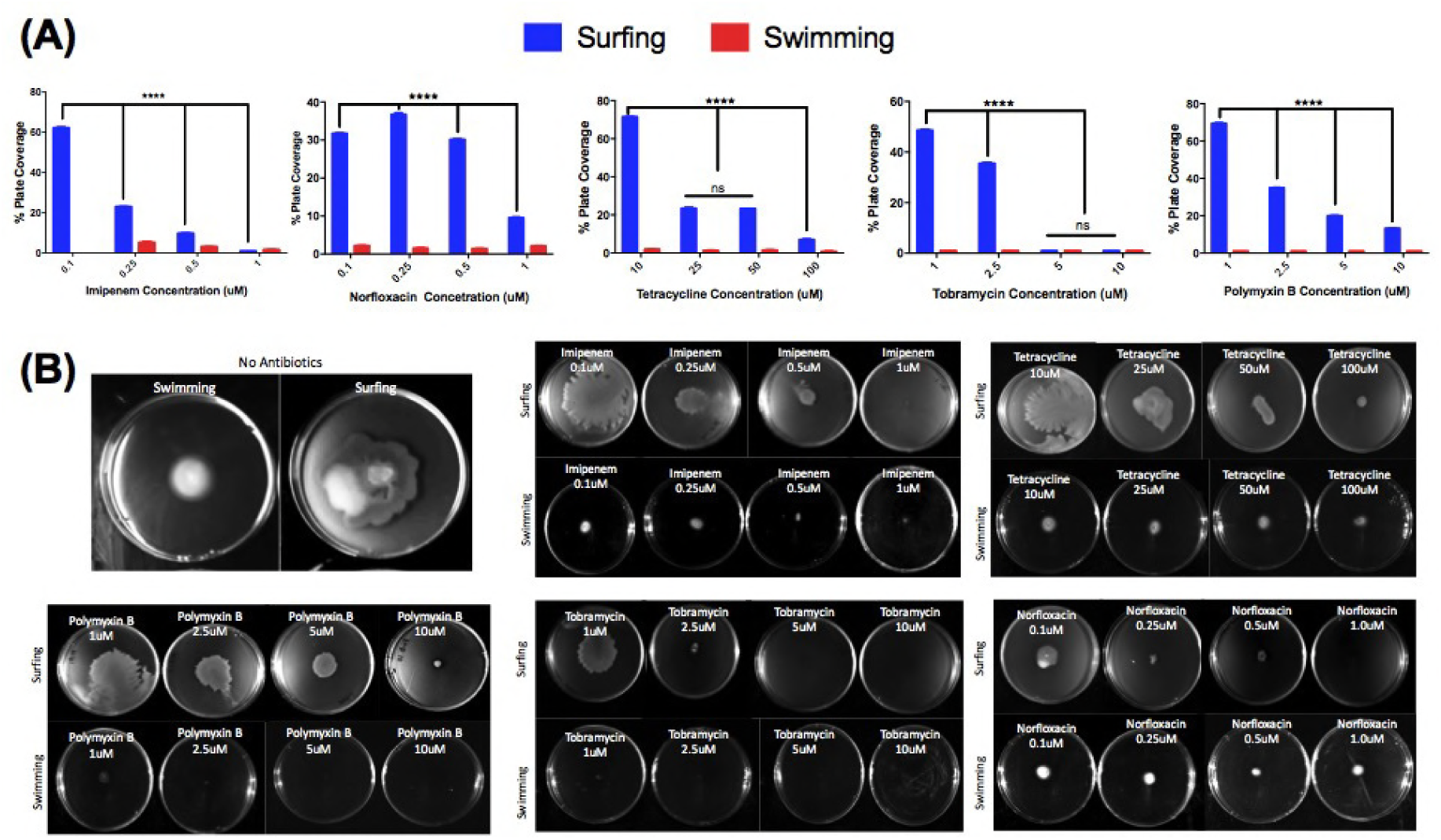
Concentration dependent inhibition of surfing motility. Surfing motility colonies of wild-type PA14 when the antibiotic at varying concentrations was incorporated into 25 mL of BM2 glucose agar containing 0.4% mucin (surfing) or no mucin (swimming). Incorporation assay results are described as the % plate coverage, relative to the control with no antibiotics, measured using Image J. Surfing colonies are represented by the blue bars and swimming by the red bars. Significance analysis between surfing and swimming was done using two-way ANOVA. * p<0.5, * * p<0.01, * * * p< 10^-3^, * * * * p<10^-4^

### Adaptive antibiotic resistance was not due to the presence of mucin alone

To show that the observed resistances were attributable to the surfing adaptation rather than the presence of mucin, we examined the effect of mucin on the broth dilution MIC of *P. aeruginosa* PA14 (Table 1). In liquid SCFM medium, the MIC values for most antibiotics, with and without mucin, remained fairly similar (no difference or a 2-fold difference). Discrepancies occurred for amikacin for which mucin increased the MIC by 4-fold, and colistin where mucin conditions resulted in an 8-fold higher MIC. This might be a result of association between the negatively charged mucin and these polycationic antibiotics. Conversely, against tetracycline mucin actually increased susceptibility by 4-fold. Overall these data suggested that the observed resistance (Fig 1, 2) was likely due to adaptation accompanying surfing motility rather than the presence per se of mucin. For this reason, we investigated these adaptive changes in greater detail.

**Table 1.** Influence of mucin on MIC. Liquid MIC results done in SCFM with 0.4% mucin and without mucin grown at 37°C overnight with an inoculums size of 2-7x10^5^cells (n=3-5).

| | MIC (μM) | |
| --- | --- | --- |
|  | + mucin | - mucin |
| Gentamicin | 4 | 1 |
| Tobramycin | 2 | 2 |
| Amikacin | 16 | 4 |
| Imipenem | 0.325 | 0.625 |
| Meropenem | 0.125 | 0.125 |
| Ceftazidime | 31.25 | 31.25 |
| Aztreonam | 4 | 8 |
| Piperacillin | 4 | 4 |
| Erythromycin | 500 | 250 |
| Clarithromycin | 2000 | 2000 |
| Polymyxin B | 16 | 16 |
| Colistin | 16 | 2 |
| Norfloxacin | 16 | 16 |
| Ciprofloxacin | 1 | 0.5 |
| Trimethoprim | 128 | 128 |
| Tetracycline | 64 | 256 |
| Chloramphenicol | 32 | 16 |

### Surfing-mediated antibiotic resistance is associated with multiple resistome genes

RNA-Seq data (NCBI GEO Accession: GSE110044) revealed that surfing is an adaptation that strongly affected gene expression. RNA-Seq was performed on two different regions of surfing colonies namely the thick white edge and blue-green centre compared to swimming cells grown in SCFM medium without mucin. In total, there were 1,467 genes dysregulated in the edge and 2,078 genes in the centre, with 816 genes commonly dysregulated between the two. To examine the possibility that adaptive resistance during surfing motility was due to the dysregulation of genes that influence resistance, literature searches were conducted. This revealed 119 genes that when mutated led to increased susceptibility (intrinsic resistance genes) and 252 genes that when mutated mediated antibiotic resistance; collectively these form the resistomes for various antibiotics (16–22). Among the resistome genes, 65 were identified, through RNA-Seq gene expression data from surfing cells, that matched the direction of dysregulation of expression levels expected if they were to have a potential role in surfing mediated resistance. Available transposon mutants of these 65 resistome genes were tested for changes in susceptibility to certain antibiotics.

Table 2 shows the resistome genes dysregulated in the edge and/or centre for which transposon mutant showed a change in susceptibility to at least one of the 5 tested antibiotics based on an initial disk diffusion assay. Several of these genes showed a change in susceptibility to more than one antibiotic, possibly illustrating a contribution to broad-spectrum resistance. The mean zone of inhibition measurements are presented in Table S1. Five of the tested mutants, Δ*recG*, Δ*ddaH*, Δ*armR* Δ*nalC*, and ΔPA3667, were similarly dysregulated in the centre and edge of a surfing colony, with *recG* and *ddaH* both up-regulated and *armR, nalC*, and *PA3667* down-regulated. Complements of selected resistome mutants showed that this broad-spectrum effect could be significantly reversed either partially, completely or excessively (Table 3), and that overexpression of some of these genes also revealed a change in susceptibility to other antibiotics as shown in Table 3. RT-qPCR data (Table S2) verified the direction of dysregulation shown in the RNA-Seq data for selected resistome genes.

**Table 2.** Resistome genes and their corresponding changes in antibiotic susceptibility relative to wild-type when mutated. This included 8 resistome genes similarly regulated in both the centre and edge. A further 10 resistome genes dysregulated only at the edge of a surfing colony were affected in such a way as to influence resistance or susceptibility. Twenty resistome genes, dysregulated only in the centre of a surfing colony, were affected in such a way as to influence antibiotic resistance or susceptibility.

| Gene | Gene Function (40) | RNA-Seq<br>Fold Change |  | Antibiotic Susceptibility of<br>Mutant Relative to WT |
| --- | --- | --- | --- | --- |
|  |  | Centre | Edge |  |
| *recG* | ATP-dependent DNA helicase | 1.9 | 2.1 | TET<sup>S</sup>, PXB<sup>S</sup>, TOB<sup>S</sup>, NFX<sup>S</sup> |
| *ddaH* | Dimethylarginine<br>dimethylaminohydrolase | 4.9 | 3.4 | IMI<sup>S</sup>, TET<sup>S</sup> |
| *armR* | Anti-repressor for MexR | -3.2 | -5.1 | IMI<sup>R</sup>, TET<sup>R</sup>, PXB<sup>R</sup>, TOB<sup>R</sup>, NFX<sup>R</sup> |
| *nalC* | Transcriptional regulator | -5.3 | -2.7 | PXB<sup>R</sup>, TOB<sup>R</sup>, NFX<sup>R</sup> |
| PA3667 | Probable pyridoxal-phosphate<br>dependent enzyme | -1.7 | -2.5 | TET<sup>R</sup> |
| PA5130 | Conserved hypothetical protein | NC | 2.4 | IMI<sup>S</sup>, TET<sup>S</sup>, PXB<sup>S</sup>, TOB<sup>S</sup>, NFX<sup>S</sup> |
| *cycH* | Cytochrome c-type biogenesis protein | NC | 2.2 | PXB<sup>S</sup>, TOB<sup>S</sup> |
| *clpS* | ATP-dependent Clp protease adaptor<br>protein | NC | -2.3 | IMI<sup>R</sup>, PXB<sup>R</sup>, TOB<sup>R</sup>, NFX<sup>R</sup> |
| PA3576 | Hypothetical protein | NC | -2.9 | TET<sup>R</sup>, TOB<sup>R</sup>, NFX<sup>R</sup> |
| PA2047 | Probable transcriptional regulator | NC | -2.0 | TOB<sup>R</sup>, NFX<sup>R</sup> |
| PA4781 | Cyclic di-GMP phosphodiesterase | NC | -2.9 | TOB<sup>R</sup>, NFX<sup>R</sup> |
| PA1513 | Hypothetical protein | NC | -3.0 | TET<sup>R</sup> |
| PA2566 | Conserved hypothetical protein | NC | -5.0 | NFX<sup>R</sup> |
| PA1348 | Hypothetical protein | NC | -3.4 | IMI<sup>R</sup>, NFX<sup>R</sup> |
| PA2571 | Probable two-component sensor | NC | -2.7 | TOB<sup>R</sup> |
| PA4766 | Conserved hypothetical protein | NC | -2.3 | TOB<sup>R</sup> |
| PA3233 | Hypothetical protein | 2.5 | NC | NOR<sup>S</sup> |
| PA4292 | Probable phosphate transporter | -6.7 | NC | IMI<sup>R</sup>, PXB<sup>R</sup>, TOB<sup>R</sup>, NFX<sup>R</sup> |
| PA1428 | Conserved hypothetical protein | -3.4 | NC | TOB<sup>R</sup>, NFX<sup>R</sup> |
| *ccoO1* | Cytochrome c oxidase, cbb3-type, CcoO subunit | -3.5 | NC | TOB<sup>R</sup>, NFX<sup>R</sup> |
| *atpB* | ATP synthase A chain | -2.1 | NC | TOB<sup>R</sup>, NFX<sup>R</sup> |
| *nuoB* | NADH dehydrogenase I chain B | -2.8 | NC | IMI<sup>R</sup>, PXB<sup>R</sup>, TOB<sup>R</sup>, NFX<sup>R</sup> |
| PA4429 | Probable cytochrome c1 precursor | -2.3 | NC | PXB<sup>R</sup>, TOB<sup>R</sup>, NFX<sup>R</sup> |
| *etfA* | Electron transfer flavoprotein alpha-subunit | -6.2 | NC | PXB<sup>R</sup>, TOB<sup>R</sup>, NFX<sup>R</sup> |
| *serA* | D-3-phosphoglycerate dehydrogenase | -12.4 | NC | PXB<sup>R</sup>, TOB<sup>R</sup>, NFX<sup>R</sup> |
| *ccmF* | Cytochrome C-type biogenesis protein | -2.2 | NC | PXB<sup>R</sup>, TOB<sup>R</sup>, NFX<sup>R</sup> |
| *nuoG* | NADH dehydrogenase I chain G | -2.1 | NC | TOB<sup>R</sup>, NFX<sup>R</sup> |
| *pchF* | Pyochelin synthetase | -2.2 | NC | TOB<sup>R</sup> |
| *rph* | Ribonuclease PH | -2.3 | NC | TOB<sup>R</sup> |
| *gidA* | Glucose-inhibited division protein A | -2.2 | NC | NFX<sup>R</sup> |
| *mutS* | DNA mismatch repair protein | -2.5 | NC | PXB<sup>R</sup>, TOB<sup>R</sup> |
| *thiG* | Thiamine biosynthesis protein, thiazole moiety | -2.9 | NC | IMI<sup>R</sup>, NFX<sup>R</sup> |
| *nuoF* | NADH dehydrogenase I chain F | -2.4 | NC | TOB<sup>R</sup> |
| *pckA* | Phosphoenolpyruvate carboxykinase | -2.6 | NC | TOB<sup>R</sup> |
| *braB* | Branched chain amino acid transporter | -4.2 | NC | NFX<sup>R</sup> |
| *htpX* | Heat shock protein | -2.0 | NC | NFX<sup>R</sup> |

**Table 3.**
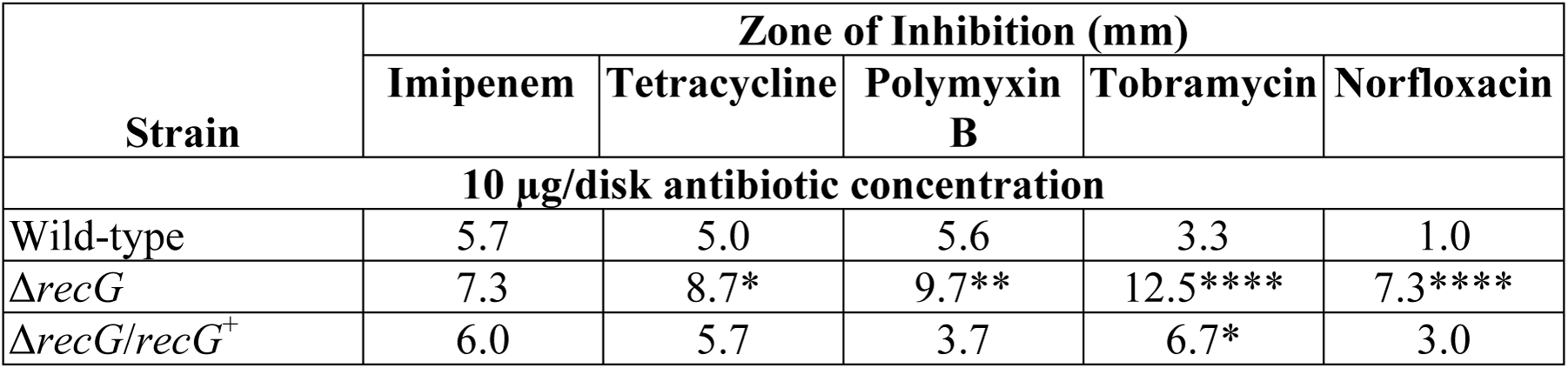
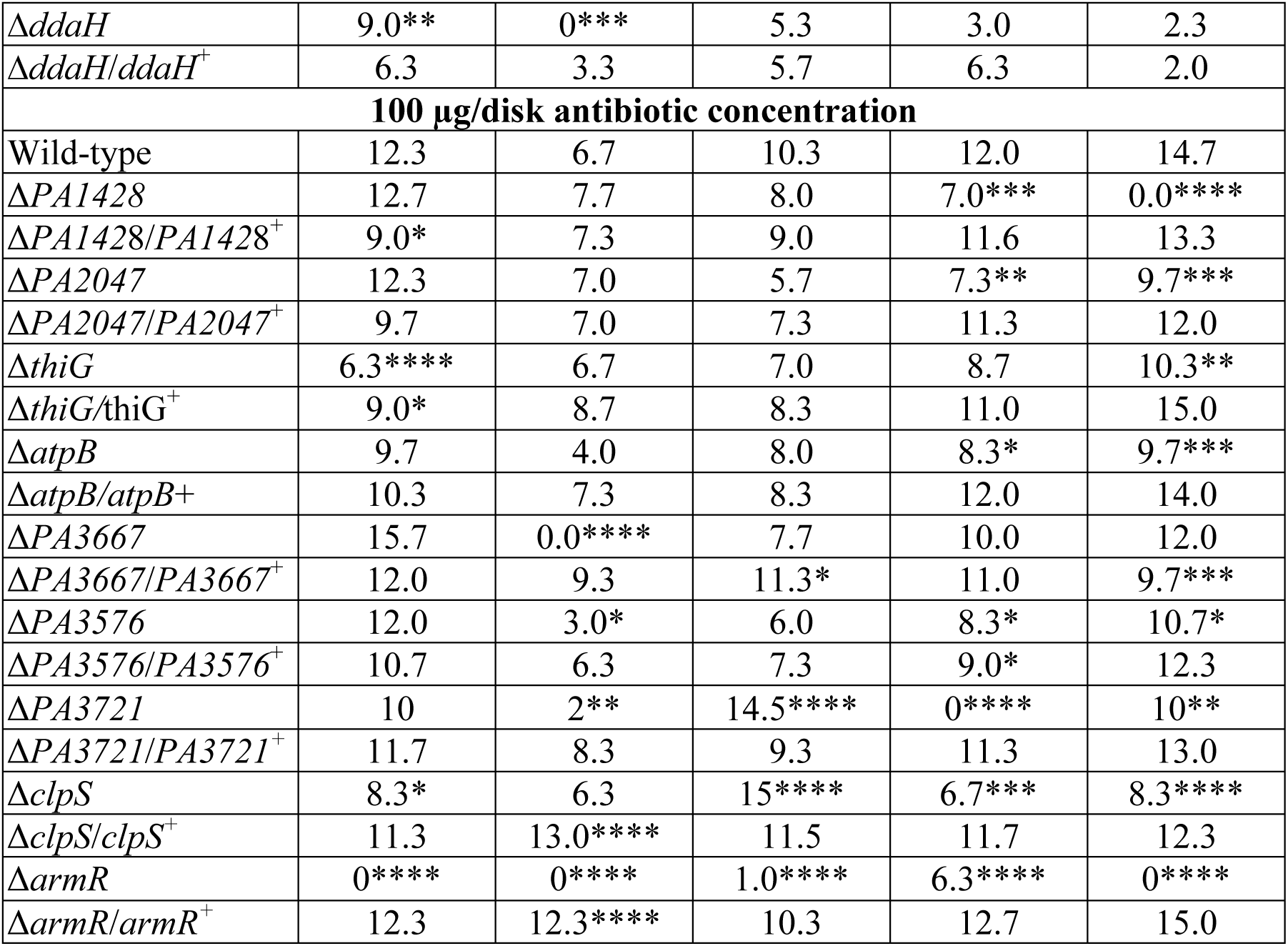
Complementation of selected resistome mutants that showed broad spectrum changes in susceptibility led to restoration of antibiotic **susceptibility**. Results show the zone of inhibition (n=3) of each mutant and its complemented equivalent against five antibiotics cf. wild-type (n=6). Mutants of up-regulated resistome genes were tested against 10ug/disk of antibiotic and down-regulated against 100ug/disk. Standard deviations ranged between 0 and 2.5mm. Statistical significance relative to wild-type was determined using two-way ANOVA. ^*^ p<0.5, ^* *^ p<0.01, ^* * *^ p> 10^-3^, ^* * **^p<10^-4^

## Discussion

*P. aeruginosa* is a highly adaptable organism that exhibits diverse lifestyles from coordinated forms of motility like swarming to community-based sessile structures like biofilms. Previously we described a new form of *P. aeruginosa* motility known as surfing under artificial cystic fibrosis-like conditions where the mucin content is high (12). Here we demonstrated that this novel form of motility is associated with multidrug adaptive resistance and is a complex adaptation influencing expression of hundreds of genes. Both disk diffusion and antibiotic incorporation assays revealed that cells undergoing surfing were significantly more resistant to multiple antibiotics compared to swimming, and the same concentrations of antibiotics that completely abolished swimming were found to be much less effective against surfing. MIC assays revealed that the observed phenomenon was dependent on surface growth associated with surfing adaption and not merely due to the presence of mucin. To explain the mechanisms behind surfing-mediated resistance, we explored the contribution of resistome genes found to be dysregulated in surfing through RNA-Seq and transposon mutant screens. In total, 36 resistome genes we identified as dysregulated under surfing conditions and that exhibited a change in susceptibility to certain antibiotics when mutated.

Swarming is another complex form of motility exhibited by *P. aeruginosa* found to be involved with major transcriptional changes (11, 23), substantially distinct from the transcriptional profile of surfing cells. Swarming has also previously been shown to be resistant to multiple antibiotics including polymyxin B, ciprofloxacin, and gentamicin, and *pvdQ* mutants influenced swarming-specific resistance (11, 23). Here surfing was also found to be associated with resistance against these same antibiotics and many others. Among the resistome genes identified in this study to be dysregulated under surfing conditions that showed contributions to adaptive antibiotic resistance, *pchF* (11, 24), *atpB, ccoO1*, and PA4429 (24) were also shown to be dysregulated under swarming conditions (11, 24). These 4 genes were found to be down-regulated in the surfing centre compared to swimming cells, and their mutant variants showed an increase in resistance to norfloxacin, tobramycin, and/or polymyxin B raising the possibility that there might be some mechanistic overlap in adaptive resistance between swarming and surfing cells as part of their complex adaptations.

Among the 36 resistome genes for which mutants showed a change in susceptibility to certain antibiotics, there were 5 that showed the same direction of dysregulation (i.e. both down or up-regulated) in both the centre and edge of a surfing colony. *RecG* and *ddaH* were both up-regulated in the surfing centre and edge, and their mutants exhibited similar reduced resistance to tetracycline. The mutant in *recG* (encoding an ATP-dependent DNA helicase) also exhibited increased susceptibility to polymyxin B, tobramycin, and norfloxacin. Tetracycline and tobramycin target protein synthesis through the 30S ribosomal submit while polymyxin B targets the cell membrane, and norfloxacin targets DNA replication. The broad-spectrum activity observed by *recG* as a resistome gene against such diverse antibiotics may arise from its regulatory nature, in that *recG* transcriptionally regulates OxyR-controlled genes in *P. putida* (25). Genes identified in the RecG regulon of *P. putida* included porins (*oprE, oprD*, PP0883) and thioredoxin reductase (trxB) involved in stress coping mechanisms (25).

There were 3 genes, *armR, nalC*, and PA3667, that were down-regulated in both regions of the surf colony. NalC is known to negatively regulate the expression of *armR*, and ArmR inhibits MexR’s DNA binding activity (26, 27). MexR negatively regulates expression of the *mexAB-oprM* operon, which encodes for a major efflux pump in *P. aeruginosa*, intrinsically involved in broad-spectrum antibiotic resistance (26). ArmR allosterically binds to MexR to alleviate its repression on the *mexAB-oprM* operon (26). Interestingly, Starr et al. (2012) revealed that a knock-out mutant of *armR* still exhibited increased expression levels of the *mexAB-oprM* operon under certain conditions (27). Here we showed that mutants in *armR* and *nalC*, which are both down-regulated in surfing, exhibited similar increases in resistance to tobramycin, norfloxacin, and polymyxin B. The observed increases in resistance to these antibiotics might be attributed in part to increased expression levels of the *mexAB-oprM* operon.

There were 11 genes dysregulated at the edge and 20 genes at the centre of a surfing colony that exhibited a change in susceptibility to at least one of the tested antibiotics when mutated compared to the wild-type. PA5130 was a conserved hypothetical protein found to be up-regulated in the surfing edge and exhibited an increased susceptibility to all 5 of the tested antibiotics when mutated. The ATP-dependent protease adapter *clpS* which was downregulated at the edge, exhibited significant increase in resistance to imipenem, polymyxin B, tobramycin, and norfloxacin. ClpS has been previously shown by our lab to contribute to antibiotic resistance, biofilm formation, and swarming motility (28). More specifically, a transposon mutant variant of *clpS* was observed to have increased resistance to β-lactams through the increased expression of of β-lactamase (28). Here it was shown that *clpS* also has an effect on resistance against imipenem, polymyxin B, tobramycin, and norfloxacin under surfing conditions.

RNA-Seq data on cells collected from the centre of a surf colony revealed 10 genes, *ccoO1, atpB, nuoB*, PA4429, *eftA, serA, ccmF, thiG, nuoF*, and *pckA*, involved in metabolism and energy production, that were down-regulated and for which mutants exhibited increased resistance to certain antibiotics. Three of these genes, *ccoO1, atpB*, and PA4429, have also been shown to be dysregulated under swarming conditions as discussed previously (11, 24). Mutants for these 10 metabolic genes that were down-regulated at the centre of surfing colonies showed an increased resistance to norfloxacin and/or tobramycin. Aminoglycosides are taken up by energy dependent mechanisms (29), and reduced metabolic activities have previously been shown in *P. aeruginosa* biofilms to contribute to resistance to tobramycin (30). Although norfloxacin has been shown to affect animal metabolism through interactions with cytochrome P450 (31), it has not been shown to affect metabolism in *P. aeruginosa.* Here we have demonstrated that reduced expression levels of certain metabolic resistome genes in the surf centre may contribute to adaptive resistance against tobramycin and/or norfloxacin.

Surfing motility is a novel form of motility that results in an lifestyle adaptation growing under conditions with high mucin. Here we demonstrate how surfing cells exhibited increased resistance attributed to large transcriptomic changes as a result of the adaptation.

## Methods and Materials

### Bacterial Strains and Complements

All screens and assays were done using the *Pseudomonas aeruginosa* UCBPP-PA14 (32) wild-type strain. All mutants used were derivatives of this strain and obtained from the PA14 Transposon Insertion Mutant Library (33). Complemented mutants were generated as follows. PCR primers listed in Table S3 were used to amplify the desired genes from strain PA14 genomic DNA. The amplified products were cloned into a TOPO vector using the Zero Blunt TOPO PCR Cloning Kit (Invitrogen). TOPO vectors containing amplified product were digested using two different enzymes, which differed depending on the gene of interest, and ligated into a pUCP18 vector containing the *lac* promoter. Vectors containing the desired genes were then transformed into their respective mutants.

### Disk diffusion assay

Disk diffusion assays were performed on synthetic cystic fibrosis media (SCFM) (34) prepared as described by Palmer et al (2007) without ammonia with 0.3% agar and 0.4% (wt/vol) mucin (surfing conditions), or with 0.3% agar without mucin (swimming conditions), or with 1.5% agar without mucin (disk diffusion control/growth control). Bacterial strains were grown in Luria broth (LB; Difco) liquid medium overnight then sub-cultured to mid-log phase (OD600=0.4-0.5). To assay motility, mid-log cultures were spotted on agar surfaces at four points around an antibiotic disk (Fig S1) impregnated with 10uL of antibiotic at concentrations indicated in Table S4. Agar plates were air-dried at 37°C for 30 min before inoculation and application of antibiotic disks. Once inoculated, plates were incubated at 37°C for 15-18 hours. The zone of inhibition surrounding the antibiotic disk was measured in millimeters using a ruler. In the case of asymmetric zones of inhibition, the average of the four sides was taken. Disk diffusion controls or growth controls were spread as lawns on plates and antibiotic disks were applied to the centre. Data collected under surfing and swimming conditions was normalized to the growth control and 2-way ANOVA was used to determine if any significant difference existed between surfing and swimming conditions. All statistical analysis was done using Graphpad Prism 7.

### Antibiotic Incorporation Assay

Incorporation assays were done on SCFM (34) using 0.3% agar with 0.4% mucin (surfing conditions) and 0.3% agar without mucin (swimming). Antibiotics were added into the agar before solidification. Once hardened, plates were air-dried for 30 minutes at 37°C before being inoculated with 1 μL of a sub-culture at an OD_600_=0.4-0.5. Plates were incubated at 37°C for 15-18 hours. Spot inoculation involved stabbing bacteria midway into the agar. The percentage of area growth on the plates was measured using ImageJ. Two-way ANOVA was used to determine if significant differences occurred between the two conditions (surfing and swimming) and between concentrations for surfing.

### Liquid Minimal Inhibitory Concentration (MIC)

Liquid MICs were conducted as described by Wiegand et al. (2008) (20. This assay was performed in liquid SCFM (34) with and without 0.4% mucin. An inoculum of 2 to 7 x10^5^ cells was used. Significant differences between MICs were taken as a 3-fold or greater change.

### RNA-Seq

PA14 was grown in liquid LB medium overnight and sub-cultured to an OD_600_=0.4-0.5. Mid-log phase cultures were used to inoculate SCFM (34) surfing and swimming plates, prepared as described above. Surfing plates were air-dried for 30 minutes before inoculation with 1 μL of culture and incubation at 37°C for 15-18 hours. Using sterile swabs, cells from the centre and edge of a surfing colony and centre of a swimming colony were collected into RNA protect bacteria reagent (Qiagen). RNA extraction was conducted using a RNeasy Mini Kit (Qiagen) according to the manufacturer’s protocol. Deoxyribonuclease treatment was performed using a TURBO DNA-free kit (Thermo Fisher) and rRNA depletion was performed using a RiboZero Bacteria Kit (Illumina). Single end cDNA libraries were constructed using a Kapa stranded Total RNA Kit (Kapa Biosystems) and libraries were sequenced on an Illumina HiSeq 2500 in rapid run mode with 100 bp reads that were base-called and de-multiplexed using built-in software on the sequencer. Fastq file quality control was performed using FastQC v0.11.5 (http://www.bioinformatics.babraham.ac.uk/projects/fastqc/) and MulitQC v0.8.dev0 (35). Fastq files were aligned to the UCBPP-PA14 genome (GenBank gene annotations) using bowtie-2 (36). Bam-sam file conversion and sorting were performed with samtools (37). Read count tables were generated with htseq-count v2.5 (38). Differential expression analysis was performed using DESeq2 (39). Fold-changes in surfing were calculated relative to swimming. Gene annotations were taken from the *Pseudomonas* Genome Database (40.

### RT-qPCR

RNA was collected as described for RNA-Seq. Reaction samples were prepared using the qScript one-step SYBR green RT-qPCR Kit (QuantaBio) with 5ng of RNA per 25μL reaction amplified in a Roche LightCycler 96. Quantification analysis was done using the comparative Ct method (41) using *rpoD* as the normalizing gene. All primers used for RT-qPCR are listed in Table S3.

## Acknowledgements

Research reported in this publication was supported by a Foundation grant from the Canadian Institutes for Health Research FDN-154287 and a grant from Cystic Fibrosis (CF) Canada, Award Number 2585. The content is solely the responsibility of the authors and does not necessarily represent the official views of the Canadian Institutes for Health Research. REWH holds a Canada Research Chair in Health and Genomics and a UBC Killam Professorship.

ES performed antibiotic screens, resistome screens, generation of complements, and manuscript writing. EEG analyzed and processed RNA-Seq data. RF prepared RNA samples for RNA-Seq. EEG and RF also contributed to manuscript writing. NL contributed to the generation of complemented strains and performed liquid MICs. Conceptualization, acquisition of funding, discussion of results and extensive editing and review of the manuscript was performed by REWH.

